# Both galactosaminogalactan and α-1,3-glucan contribute to aggregation of *Aspergillus oryzae* hyphae in liquid culture

**DOI:** 10.1101/589408

**Authors:** Ken Miyazawa, Akira Yoshimi, Motoaki Sano, Fuka Tabata, Asumi Sugahara, Shin Kasahara, Ami Koizumi, Shigekazu Yano, Tasuku Nakajima, Keietsu Abe

## Abstract

Filamentous fungi generally form aggregated hyphal pellets in liquid culture. We previously reported that α-1,3-glucan-deficient mutants of *Aspergillus nidulans* did not form hyphal pellets and their hyphae were fully dispersed, and we suggested that α-1,3-glucan functions in hyphal aggregation. Yet, *Aspergillus oryzae* α-1,3-glucan-deficient (AGΔ) mutants still form small pellets; therefore, we hypothesized that another factor responsible for forming hyphal pellets remains in these mutants. Here, we identified an extracellular matrix polysaccharide galactosaminogalactan (GAG) as such a factor. To produce a double mutant of *A. oryzae* (AG-GAGΔ), we disrupted the genes required for GAG biosynthesis in an AGΔ mutant. Hyphae of the double mutant were fully dispersed in liquid culture, suggesting that GAG is involved in hyphal aggregation in *A. oryzae*. Addition of partially purified GAG fraction to the hyphae of the AG-GAGΔ strain resulted in formation of mycelial pellets. Acetylation of the amino group in galactosamine of GAG weakened GAG aggregation, suggesting that hydrogen bond formation by this group is important for aggregation. Genome sequences suggest that α-1,3-glucan, GAG, or both are present in many filamentous fungi and thus may function in hyphal aggregation in these fungi. We also demonstrated that production of a recombinant polyesterase, CutL1, was higher in the AG-GAGΔ strain than in the wild-type and AGΔ strains. Thus, controlling hyphal aggregation factors of filamentous fungi may increase productivity in the fermentation industry.

**Importance:** Production using filamentous fungi is an important part of the fermentation industry, but hyphal aggregation in these fungi in liquid culture limits productivity compared with that of yeast or bacterial cells. We found that galactosaminogalactan and α-1,3-glucan both function in hyphal aggregation in *Aspergillus oryzae*, and that the hyphae of a double mutant deficient in both polysaccharides become fully dispersed in liquid culture. We also revealed the relative contribution of α-1,3-glucan and galactosaminogalactan to hyphal aggregation. Recombinant protein production was higher in the double mutant than in the wild-type strain. Our research provides a potential technical innovation for the fermentation industry that uses filamentous fungi, as regulation of the growth characteristics of *A*. *oryzae* in liquid culture may increase productivity.

## Introduction

The hyphae of filamentous fungi generally form aggregated pellets in liquid culture. Although filamentous fungi have been used for industrial production of enzymes and secondary metabolites for a long time (1, 2), hyphal pellet formation decreases productivity in liquid culture (3, 4). Formation of hyphal pellets might be related to a property of the cell surface (5), and elucidation of the relationship between hyphal aggregation and cell surface components, especially polysaccharides, is needed.

The fungal cell wall is essential for survival because it maintains the cell’s shape, prevents cell lysis, and protects cells from environmental stresses (6). Fungal cell walls are composed mainly of polysaccharides. In *Aspergillus* species, the cell wall is composed of α-glucan (mainly α-1,3-glucan), β-1,3/1,6-glucan, galactomannan, and chitin (6–8). Cell walls of some filamentous fungi are covered with extracellular matrix, which is composed mainly of polysaccharides, including α-glucan (α-1,3-glucan with a small amount of α-1,4-glucan), galactomannan, or galactosaminogalactan (GAG) (9, 10).

We reported that the Δ*agsB* and Δ*agsA*Δ*agsB* strains of *Aspergillus nidulans* have no α-1,3-glucan in the cell wall (11) and their hyphae are fully dispersed in liquid culture, whereas the wild-type strain forms aggregated pellets. In *Aspergillus fumigatus*, addition of α-1,3-glucanase prevents aggregation of germinating conidia (12). These findings strongly suggest that α-1,3-glucan is an adhesive factor. We disrupted the three α-1,3-glucan synthase genes in the industrial fungus *Aspergillus oryzae* (Δ*agsA*Δ*agsB*Δ*agsC*; AGΔ) and confirmed the loss of α-1,3-glucan in the cell wall of the AGΔ strain, but the strain still formed small hyphal pellets in liquid culture (13). Although the AGΔ hyphae were not fully dispersed, the strain produced more recombinant polyesterase (cutinase) CutL1 than did a wild-type strain (WT-cutL1) because of the smaller pellets of the AGΔ strain (13). We predicted that another cell wall or cell surface component is responsible for hyphal aggregation in the AGΔ strain. Identification of this factor, distinct from α-1,3-glucan, is important, because full dispersion of *A. oryzae* hyphae would enable higher cell density and increase production of commercially valuable products in liquid culture.

GAG is a hetero-polysaccharide composed of linear α-1,4-linked galactose (Gal), *N*-acetylgalactosamine (GalNAc), and galactosamine (GalN). GAG is an important pathogenetic factor in the human pathogen *A. fumigatus* (14, 15); it is involved in adherence to host cells, biofilm formation, and avoidance of immune response by masking β-1,3-glucan and chitin (9, 16). Disruption of genes encoding the transcription factors StuA and MedA significantly decreases GAG content and has led to identification of the *uge3* (UDP-glucose 4-epimerase) gene (16). Four genes (*sph3*, *gtb3*, *ega3*, and *agd3*) located near *uge3* have been identified (17). In *stuA* and *medA* gene disruptants, these five genes are downregulated, suggesting that they are co-regulated by StuA and MedA (17). GAG biosynthesis by the five encoded proteins is predicted in *A. fumigatus* (9, 18). First, the epimerase Uge3 produces UDP-galactopyranose (Gal*p*) from UDP-glucose and UDP-*N*-GalNAc from UDP-*N*-acetylglucosamine (GlcNAc) (16, 19). Second, glycosyltransferase Gtb3 polymerizes UDP-Gal*p* and UDP-GalNAc and exports the polymer from the cytoplasm. Third, deacetylase Agd3 deacetylates the synthesized GAG polymer (17). The predicted glycoside hydrolase Ega3 has yet to be characterized. Sph3 belongs to a novel glycoside hydrolase family, GH135, and is essential for GAG production (18), but its role in GAG synthesis remains unknown.

Here, we confirmed that *A. oryzae* has the GAG biosynthetic gene cluster. We disrupted *sphZ* (ortholog of *A. fumigatus sph3*) and *ugeZ* (ortholog of *uge3*) in the wild-type and AGΔ strains to produce Δ*sphZ*Δ*ugeZ* (GAGΔ) and Δ*agsA*Δ*agsB*Δ*agsC*Δ*sphZ*Δ*ugeZ* (AG-GAGΔ), respectively. In liquid culture, the hyphae of the AG-GAGΔ strain were fully dispersed, suggesting that GAG plays a role in hyphal adhesion in *A. oryzae*, along with α-1,3-glucan. Using the wild-type, AGΔ, GAGΔ, and AG-GAGΔ strains of *A. oryzae*, we characterized hyphal aggregation and discuss its mechanism in *A. oryzae*. Our findings may have wide implications because the genomes of many filamentous fungi encode enzymes required for α-1,3-glucan or GAG biosynthesis, or both (7, 17).

## MATERIALS AND METHODS

### Strains and growth media

Strains used are listed in Table 1. *Aspergillus oryzae* NS4 (*sC*^-^, *niaD*^-^) with Δ*ligD* (Δ*ligD*::*sC*, Δ*adeA*::*ptrA*) was used for all genetic manipulations (20). All *A. oryzae* strains were cultured in standard Czapek–Dox (CD) medium as described previously (11, 13). The *niaD*^-^ strains were cultured in CDE medium (CD medium containing 70 mM sodium hydrogen L(+)-glutamate monohydrate as the nitrogen source instead of sodium nitrate).

**Table 1.**
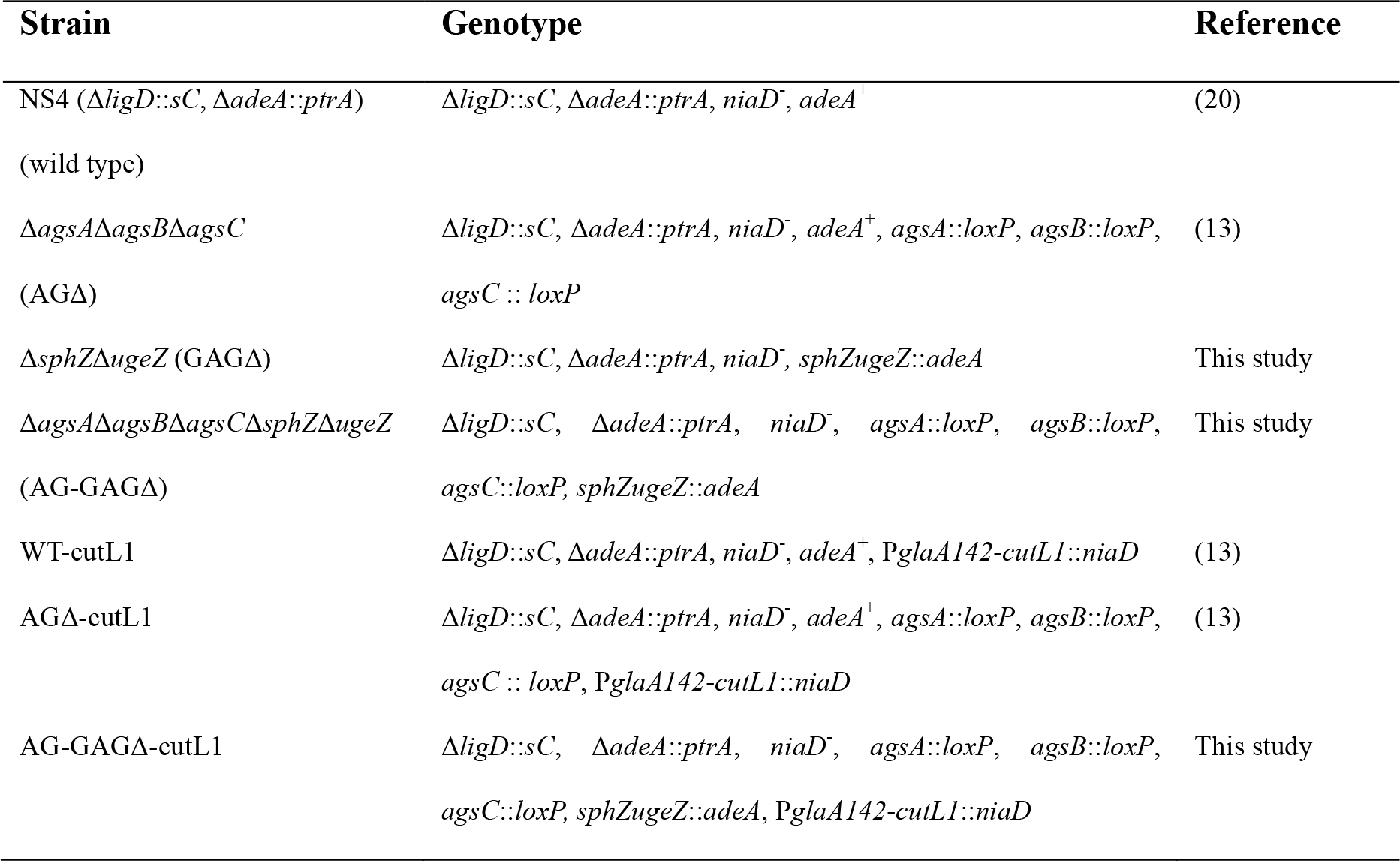
**Strains used in this study**

**Table 2.**
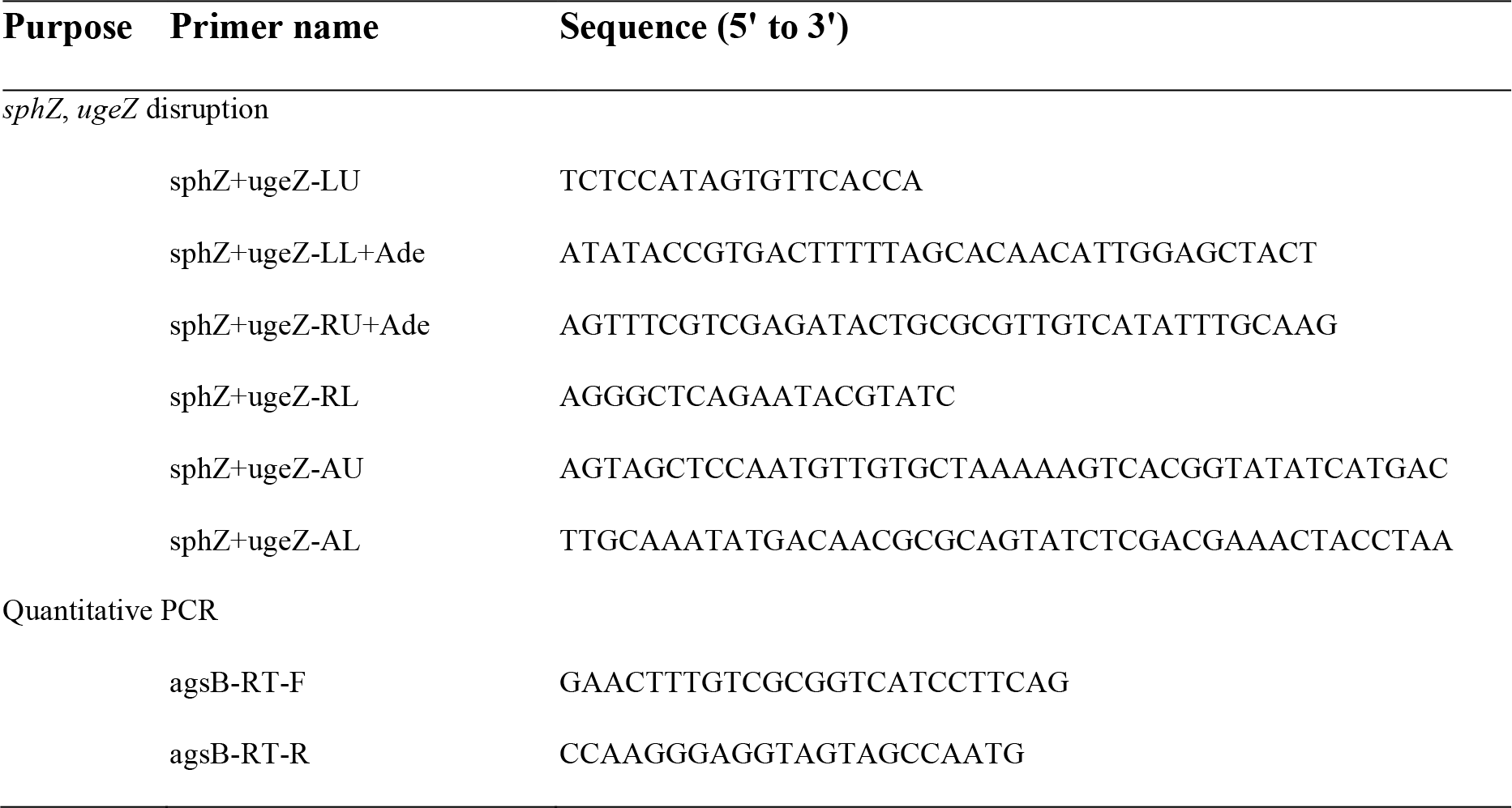
**PCR primers used in this study**

Conidia of *A*. *oryzae* used to inoculate flask cultures were isolated from cultures grown on malt medium, as described previously (13). YPD medium containing 2% peptone (Becton Dickinson and Company, Sparks, Nevada, USA), 1% yeast extract (Becton Dickinson and Company), and 2% glucose was used for flask culture to analyze growth characteristics. YPM medium containing 2% peptone, 1% yeast extract, and 2% maltose was used for flask culture to evaluate production of recombinant cutL1.

### Construction of dual *sphZ ugeZ* gene disruptant in *A. oryzae*

Fragments containing the 3′ non-coding regions of *ugeZ* (amplicon 1) and *sphZ* (amplicon 2) derived from *A. oryzae* genomic DNA, and the *adeA* gene (amplicon 3) from the TOPO-2.1-adeA plasmid (13), were amplified by PCR. Amplicon 1 was amplified with the primers sphZ+ugeZ-LU and sphZ+ugeZ-LL+ade, amplicon 2 with the primers sphZ+ugeZ-RU+ade and sphZ+ugeZ-RL, and amplicon 3 with the primers sphZ+ugeZ-AU and sphZ+ugeZ-AL. The primers sphZ+ugeZ-LL+ade, sphZ+ugeZ-AU, and sphZ+ugeZ-AL were chimeric; each contained a reverse-complement sequence for PCR fusion. The PCR products were gel-purified and used as substrates for the second round of PCR with the primers sphZ+ugeZ-LU and sphZ+ugeZ-RL to fuse the three fragments (Fig. S1A). The resulting major PCR product was gel-purified and used to transform *A. oryzae* wild-type and AGΔ strains (Fig. S1B). Disruption of the *sphZ* and *ugeZ* genes was confirmed by Southern blot analysis (Fig. S1C).

### Analysis of the growth characteristics of *A. oryzae* in liquid culture

Conidia (final concentration, 1 × 10^5^/mL) of the wild-type, AGΔ, GAGΔ, and AG-GAGΔ strains were inoculated into 50 mL of YPD medium in 200-mL Erlenmeyer flasks and rotated at 120 rpm at 30°C for 24 h. The mean diameter of the hyphal pellets was determined as described previously (13).

### Scanning electron microscopy

Conidia (final concentration, 1 × 10^5^/mL) of the wild-type, AGΔ, GAGΔ, and AG-GAGΔ *A. oryzae* strains were inoculated and grown as above. The culture broths were filtered through Miracloth (Merck Millipore, Darmstadt, Germany). The mycelia were washed with water twice, dehydrated with tert-butanol, lyophilized, and coated with platinum–vanadium. Mycelia were observed under a Hitachi SU8000 scanning electron microscope (Hitachi, Tokyo, Japan) at an accelerating voltage of 3 kV.

### Assay of cell wall susceptibility to Lysing Enzymes

Susceptibility of the fungal cell wall to Lysing Enzymes (LE), a commercial preparation containing β-1,3-glucanase and chitinase (Sigma, St. Louis, MO, USA), was assayed as described previously (11). Washed 1-day-old mycelia of the wild-type, AGΔ, GAGΔ, and AG-GAGΔ strains (30 mg fresh weight) grown in CDE medium at 30°C were suspended in 1 mL of 0.8 M NaCl in sodium phosphate buffer (10 mM, pH 6.0) containing 10 mg/mL LE and incubated for 1, 2, or 4 h at 30°C. The number of protoplasts generated from the mycelia was counted with a hemocytometer (A106, SLGC, Tokyo, Japan).

### Assay for growth inhibition by Congo Red

Sensitivity of the wild-type, AGΔ, GAGΔ, and AG-GAGΔ strains to Congo Red was evaluated using our previously described method (11), with a minor modification. Briefly, conidial suspensions of each strain (1.0 × 10^4^ cells) were spotted on the centers of CDE plates containing Congo Red (10, 20, 40, 80, or 120 µg/mL) and incubated at 30°C for 3 days. The dose response was determined by plotting the mean diameters of the colonies on media with Congo Red as a percentage of those on control medium. Each experiment was performed in quadruplicate.

### Fractionation of cell wall components and quantification of carbohydrate composition

Conidia (final concentration, 1.0 × 10^5^/mL) of the wild-type, AGΔ, GAGΔ, and AG-GAGΔ strains were inoculated into 200 mL of YPD medium in 500 mL-Erlenmeyer flasks and rotated at 120 rpm at 30°C. Mycelia were collected by filtration through Miracloth, washed twice with 20 mL of water, and lyophilized. Mycelia were pulverized with a MM400 bench-top mixer mill (Retsch, Haan, Germany), and the resulting powder (1 g) was suspended in 40 mL of 0.1 M sodium phosphate buffer (pH 7.0). Cell wall components were fractionated by hot-water and alkali treatment (13), and the fractionation resulted in hot-water-soluble (HW), alkali-soluble (AS), and alkali-insoluble (AI) fractions. The AS fraction was further separated into a fraction soluble in water at neutral pH (AS1) and an insoluble fraction (AS2). The carbohydrate composition of the fractions was quantified as described previously (11). Briefly, 10 mg of each cell wall fraction was hydrolyzed with sulfuric acid and then neutralized with barium sulfate. The carbohydrate composition of the hydrolysate was determined using high-performance anion-exchange chromatography (HPAEC). For GalN quantification, the carbohydrate composition of sulfuric acid–hydrolyzed HW fractions (50 mg each) was quantified.

### Purification of GAG from culture supernatant of the AGΔ strain by fractional precipitation with ethanol

Conidia (final concentration, 1.0 × 10^6^/mL) of the AGΔ or AG-GAGΔ strain (negative control) were inoculated into 3 flasks, each containing 1 L of modified Brian medium (14), and rotated at 160 rpm at 30°C for 72 h. The mycelia were removed by filtration through Miracloth. The supernatants were combined and concentrated to 1 L by evaporation, dialyzed against water at 4°C, and concentrated again to 1 L; then 20 g of NaOH was added (final concentration, 0.5 M) at 4°C with stirring. The mixture was centrifuged at 3000 ×*g* at 4°C for 10 min and a pellet was obtained (referred to hereafter as the 0 vol). EtOH fraction. EtOH (0.5 L) was added to the supernatant and the mixture was incubated for 5 h at 4°C with stirring, then centrifuged at 3000 ×*g* at 4°C for 10 min, and a pellet (0.5 vol. EtOH fraction) was obtained. These procedures were repeated to obtain 1, 1.5, 2, and 2.5 vol. EtOH fractions. Each fraction was neutralized with 3 M HCl, dialyzed against water, and freeze-dried. The carbohydrate composition of each fraction was determined as above. For mycelial aggregation assay, each freeze-dried EtOH fraction from the AGΔ strain (2 mg) and the 1.5 vol. EtOH fraction from the AG-GAGΔ strain was dissolved in 1 mL of 0.1 M HCl and vortexed for 10 min.

### Conidial and mycelial aggregation assay

A modified method of Fontaine et al. (12) was used. Conidia (5 × 10^5^) were inoculated into 500 µL of CDE liquid medium containing 0.05% Tween 20 in a 48-well plate and agitated at 1200 rpm with a microplate mixer (NS-P; As One, Osaka, Japan) at 30°C for 3, 6, or 9 h. Conidial aggregates were then examined under a stereomicroscope (M125; Leica Microsystems, Wetzlar, Germany). Mycelial aggregation in the presence of GAG was evaluated as follows. Conidia (final concentration, 1.0 × 10^7^/mL) of the AG-GAGΔ strain were inoculated into 50 mL of YPD medium and rotated at 120 rpm at 30°C for 9 h. The mycelia were collected by filtration through Miracloth and washed twice with water. The mycelia (wet weight, 500 mg) were resuspended in 10 mL of PBS, and the suspension (25 µL) was added into a mixture of 400 µL of water, 50 µL of 1 M sodium phosphate buffer (pH 7.0), and 25 µL of the EtOH fraction (from AGΔ) or the mock fraction (from AG-GAGΔ). Aggregates were examined under a stereomicroscope after 1 h.

To evaluate the effect of pH on mycelial aggregation by GAG, the mycelial suspension (25 µL) was added into a mixture of 450 µL of buffer (final concentration, 100 mM) and 25 µL of the 1.5 vol. EtOH fraction. The following buffers were used: pH 4.0–5.0, sodium acetate; pH 6.0–7.0, sodium phosphate; pH 8.0, Tricine-NaOH. Aggregates were examined at 1 h.

To examine the effect of inhibiting hydrogen bond formation, the mycelial suspension (25 µL) was added into a mixture of 450 µL of 100 mM sodium phosphate buffer (pH 7.0) and 0, 1, 2, 4, or 8 M urea, and 25 µL of the 1.5 vol. EtOH fraction.

### Acetylation of the amino group of galactosaminogalactan

The 1.5 vol. EtOH fraction from the AGΔ (5 mg) was dissolved in ice cold 0.5 M NaOH (800 µL), neutralized with ice cold 2 M HCl, and then added to 4 mL of 50 mM sodium acetate. Then, methanol (4 mL) and acetic anhydrate (10 mg) were added and the mixture was stirred at room temperature for 24 h. The sample was then evaporated, washed three times with methanol, dialyzed against water, and freeze-dried. The procedure was then repeated.

### Visualization of α-1,3-glucan, GAG, and hydrophobin in the cell wall

Germinating conidia cultured in a 48-well plate were dropped onto a glass slide, washed twice with PBS, and fixed with 4% (w/v) paraformaldehyde for 10 min. Samples were washed twice with 50 mM potassium phosphate buffer (pH 6.5) and stained at room temperature for 2 h with Alexa Fluor 647–conjugated soybean agglutinin (SBA; 100 µg/mL) (Invitrogen) and α-1,3-glucanase-α-1,3-glucan–binding domain fused with GFP (AGBD-GFP; 100 µg/mL) (21) in 50 mM phosphate buffer (pH 6.5). After being washed with the same buffer, the samples were imaged under a FluoView FV1000 confocal laser-scanning microscope (Olympus, Tokyo, Japan). For hydrophobin staining, fixed cells were washed twice with PBS and incubated at 30°C for 2 h with a rabbit polyclonal antibody against RolA (T.K. Craft, Gunma, Japan) diluted 1:200 in PBS. Cells were then washed three times with PBS, and a drop of PBS containing secondary antibody (anti-rabbit IgG antibody–Alexa Fluor 568 conjugate; Invitrogen) was added. The sample was incubated at room temperature for 1 h, washed as above, and imaged under the confocal laser-scanning microscope.

### Quantification of CutL1 production

A *cutL1*-overexpressing strain (AG-GAGΔ-cutL1) was constructed as described previously (13) with the pNGA-gla-Cut plasmid (22). Integration of a single copy of the *cutL1-*overexpression construct at the *niaD* locus was confirmed by Southern blot analysis (Fig. S2). Enzyme production in the mutants was evaluated as described previously (13), with some modifications. Briefly, conidia (final concentration, 1 × 10^4^/mL) of the WT-cutL1, AGΔ-cutL1, and AG-GAGΔ-cutL1 strains were inoculated into 50 mL of YPM medium and rotated at 100 rpm at 30°C for 24 h. The culture broth was filtered through Miracloth. Mycelial cells were dried at 70°C for 24 h and weighed. Proteins were precipitated from an aliquot of the filtrate (400 µL) with 100% (w/v) trichloroacetic acid (200 µL), separated by SDS-PAGE, and stained with Coomassie Brilliant Blue. ImageJ software was used to quantify the amount of CutL1 in the broth; purified CutL1 was used for calibration.

### ^13^C NMR analysis of cell wall fractions

^13^C NMR analysis was performed as described previously (23). The AS2 fractions from the wild-type and GAGΔ strains were dissolved in 1 M NaOH/D_2_O. Me_2_SO-d_6_ (deuterated dimethyl sulfoxide; 5 µL) was added to each sample. ^13^C NMR spectra were obtained using a JNM-ECX400P spectrometer (JEOL, Tokyo, Japan) at 400 MHz, 35°C (72,000 scans).

### RNA purification, reverse transcription, and quantitative PCR

Total RNA was extracted using Sepasol-RNA I Super according to the manufacturer’s instructions (Nakarai Tesque, Kyoto, Japan). Total RNA (2 µg) was reverse transcribed and cDNA was amplified using a High-Capacity cDNA Reverse Transcription Kit according to the manufacturer’s instructions (Thermo Fisher Scientific, Waltham, MA, USA). Quantitative PCR was performed with the AoagsB-RT-F and AoagsB-RT-R primers using KOD SYBR qPCR Mix (Toyobo Co., Ltd., Osaka, Japan).

### Statistical analysis

Tukey’s test was used for the comparison of multiple samples.

## Results

### *Aspergillus oryzae* has a GAG biosynthetic gene cluster

In *A. fumigatus*, GAG biosynthesis is regulated by a cluster of five genes, and this cluster is conserved in a wide range of filamentous fungi (17).To check whether *A. oryzae* possesses the GAG gene cluster, we used a BLAST search (https://blast.ncbi.nlm.nih.gov/Blast.cgi). We found that all five GAG biosynthetic genes, orthologous to *A. fumigatus uge3*, *sph3*, *ega3*, *agd3*, and *gtb3* (Fig. 1A), are conserved in *A. oryzae*: *ugeZ*, *sphZ*, *egaZ*, *agdZ*, and *gtbZ* (Fig. 1B). *Aspergillus oryzae* UgeZ had motifs conserved among group 2 epimerases (19), SphZ contained a spherulin 4 conserved region (18), and AgdZ had the conserved motifs of the carbohydrate esterase family 4 (17). These findings indicated that *A*. *oryzae*, similar to *A*. *fumigatus*, can produce GAG.

**FIG 1.**
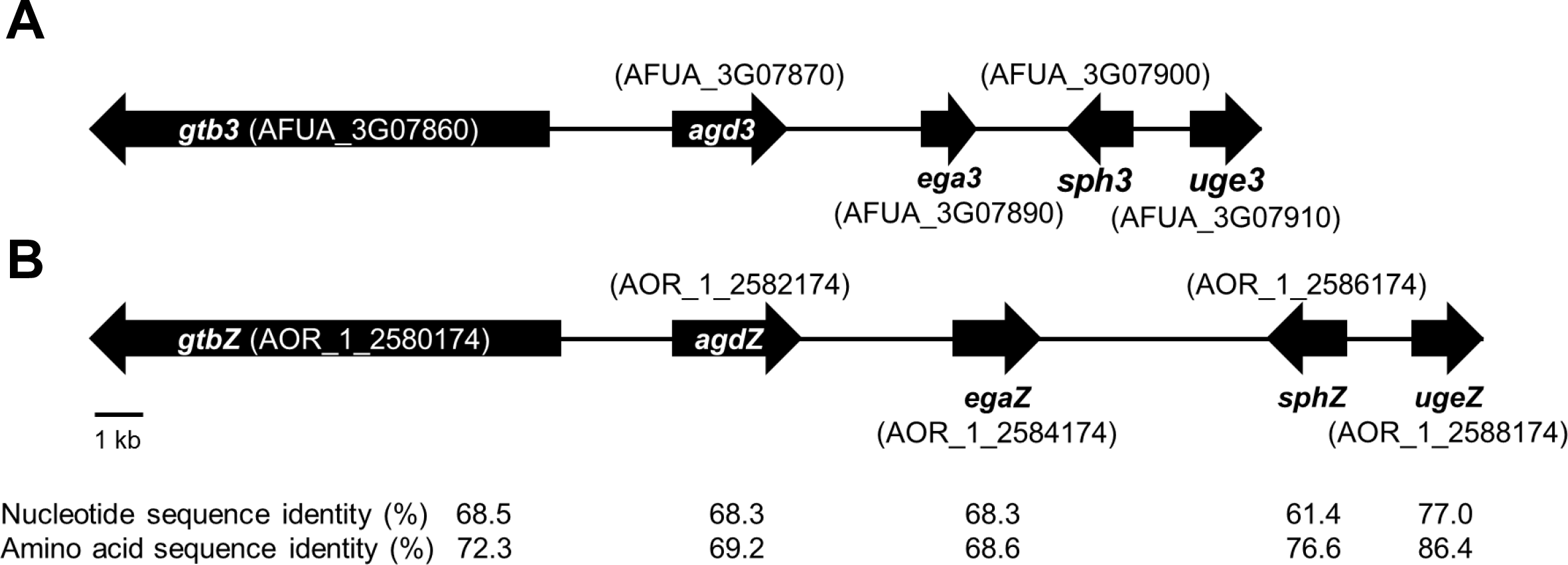
GAG biosynthetic cluster in (A) *Aspergillus fumigatus* and (B) *Aspergillus oryzae.* The cluster of *A*. *oryzae* was predicted from the sequence of the cluster of *A*. *fumigatus* by using a BLAST search.

### Hyphae of the AG-GAGΔ strain are completely dispersed in liquid culture

Because disruption of the *sph3* and *uge3* genes leads to a loss of GAG in *A. fumigatus* (18, 19), we disrupted *sphZ* and *ugeZ* in *A. oryzae* in the genetic background of the wild-type and AGΔ strains, and we obtained the GAGΔ and AG-GAGΔ strains, respectively. The wild-type, AGΔ, AG-GAGΔ, and GAGΔ strains showed almost the same mycelial growth and conidiation on CD agar plates after 5 days at 30°C (Fig. S3). When grown in YPD liquid medium at 30°C for 24 h, the wild-type strain formed significantly larger hyphal pellets (3.7 ± 0.2 mm in diameter) than did the AGΔ strain (2.7 ± 0.3 mm; Fig. 2A, B), in good agreement with our previous results (13). The hyphae of the AG-GAGΔ strain were completely dispersed, and the GAGΔ strain formed significantly larger hyphal pellets (6.2 ± 0.0 mm) than did the wild-type strain (Fig. 2A, B). These results strongly suggest that, in addition to α-1,3-glucan, GAG has a role in hyphal adhesion in *A. oryzae* and that the defect in both AG and GAG biosynthetic genes is required for full dispersion of *A. oryzae* hyphae.

**FIG 2.**
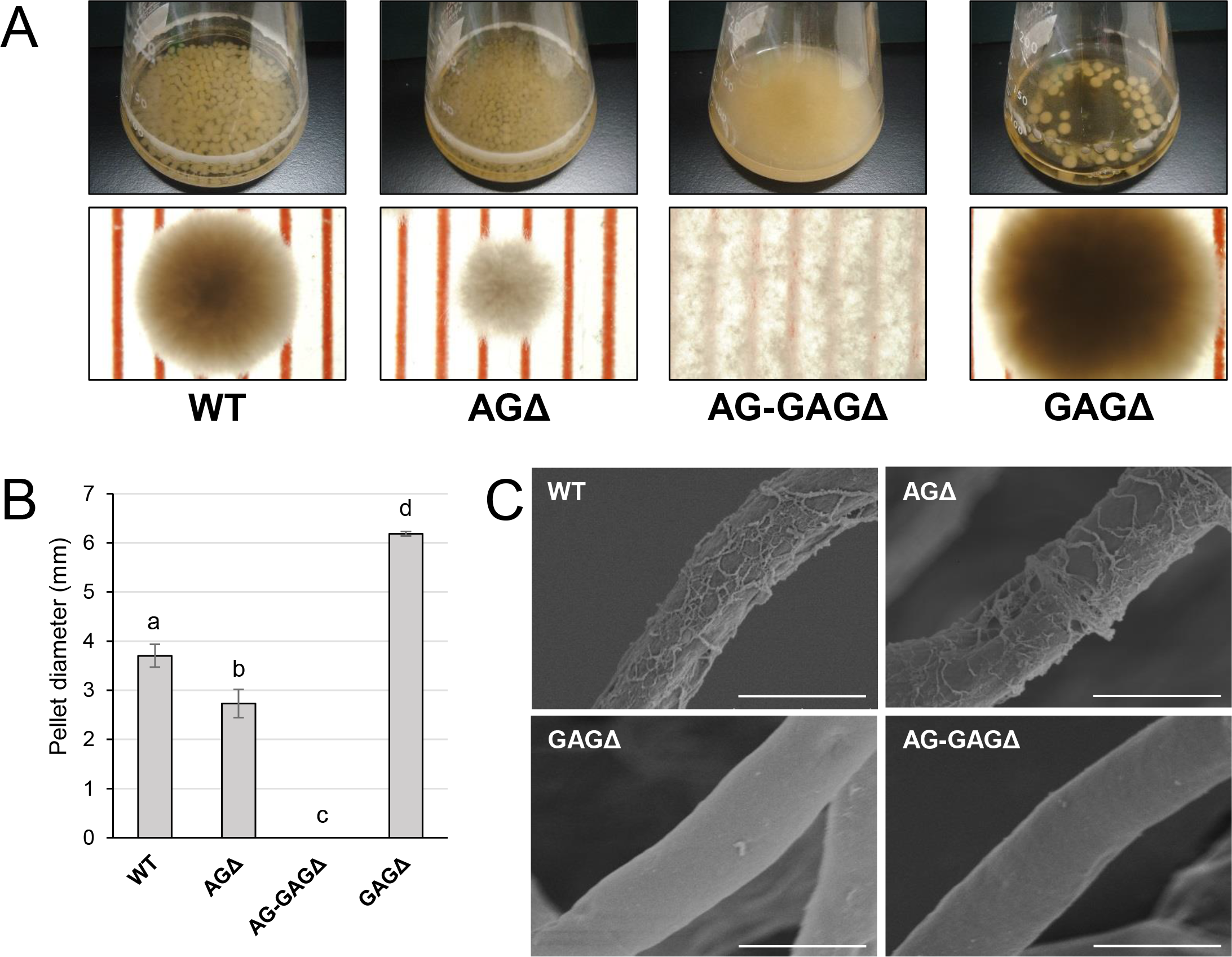
Phenotypes of *Aspergillus oryzae* Δ*agsA*Δ*agsB*Δ*agsC*Δ*sphZ*Δ*ugeZ* (AG-GAGΔ) and Δ*sphZ*Δ*ugeZ* (GAGΔ) strains in liquid culture. (A) The wild-type (WT), Δ*agsA*Δ*agsB*Δ*agsC* (AGΔ), AG-GAGΔ, and GAGΔ strains were cultured in Erlenmeyer flasks (upper row), and images of hyphal pellets were taken under a stereomicroscope (bottom row; scale, 1 mm) at 24 h of culture. (B) The mean diameter of hyphal pellets was determined by measuring 10 randomly selected pellets per replicate under a stereomicroscope. Error bars represent standard deviations calculated from three replicates. (C) Morphology of each strain was examined under a scanning electron microscope. Scale bars, 5 µm.

Scanning electron microscopy revealed that the surface of *A. fumigatus* hyphae has GAG-dependent decorations in liquid culture, which are lost in the *sph3*, *uge3*, and *agd3* gene disruptants (17–19). We investigated whether the hyphae of *A. oryzae* GAGΔ and AG-GAGΔ strains lack such decorations. As expected, we observed fibrous decorations on the hyphal cells of the wild-type and AGΔ *A. oryzae* strains (Fig. 2C), but the hyphae of the GAGΔ and AG-GAGΔ strains had smooth surfaces (Fig. 2C). These results suggest that the fibrous decorations on the cell surface are attributable to the presence of the GAG biosynthetic gene cluster in *A. oryzae*.

We used three approaches to analyze why the GAGΔ strain formed larger hyphal pellets in liquid culture: (1) HPAEC-pulsed amperometric detection analysis of cell wall components in alkali-soluble fractions showed no significant difference in the amount of glucose in the AS2 fractions between the wild-type and GAGΔ strains (Fig. S4A); (2) Expression of the *agsB* gene, which encodes the main α-1,3-glucan synthase of *A*. *oryzae* (24), was slightly lower in the GAGΔ strain than in the wild-type strain at 6 h of culture, but it was slightly higher at 24 h (Fig. S4B); and (3) ^13^C NMR analysis of the AS2 fraction showed that the main component was α-1,3-glucan in both strains (Fig. S4C). The reason why the GAGΔ strain formed larger aggregated pellets remains unclear from the results of our experiments.

### Disruptants of AG and GAG biosynthetic genes are sensitive to Lysing Enzymes and Congo Red

To investigate the consequences of cell wall alteration caused by the loss of GAG, we assessed the susceptibility of the wild-type, AGΔ, AG-GAGΔ, and GAGΔ strains to LE and CR. The concentrations of protoplasts formed from hyphae tended to be higher for the AGΔ strain than for the wild-type strain after 2 and 4 h of treatment with LE (0.05 < *p* < 0.1; Fig. 3A). In contrast, protoplast concentration was significantly higher for the AG-GAGΔ strain than for the wild-type and AGΔ strains at each time point (Fig. 3A). Protoplast concentration was also significantly higher for the GAGΔ strain than for the wild-type and AGΔ strains after 1 and 4 h (Fig. 3A), but it was significantly lower than for the AG-GAGΔ strain after 4 h (Fig. 3A). All three mutant strains were significantly more sensitive to CR than the wild type: the AG-GAGΔ strain was most sensitive, and the AGΔ and GAGΔ strains showed similar sensitivity (Fig. 3B). These data revealed that AG and GAG additively contribute to cell wall protection from cell wall–degrading enzymes and environmental chemicals.

**FIG 3.**
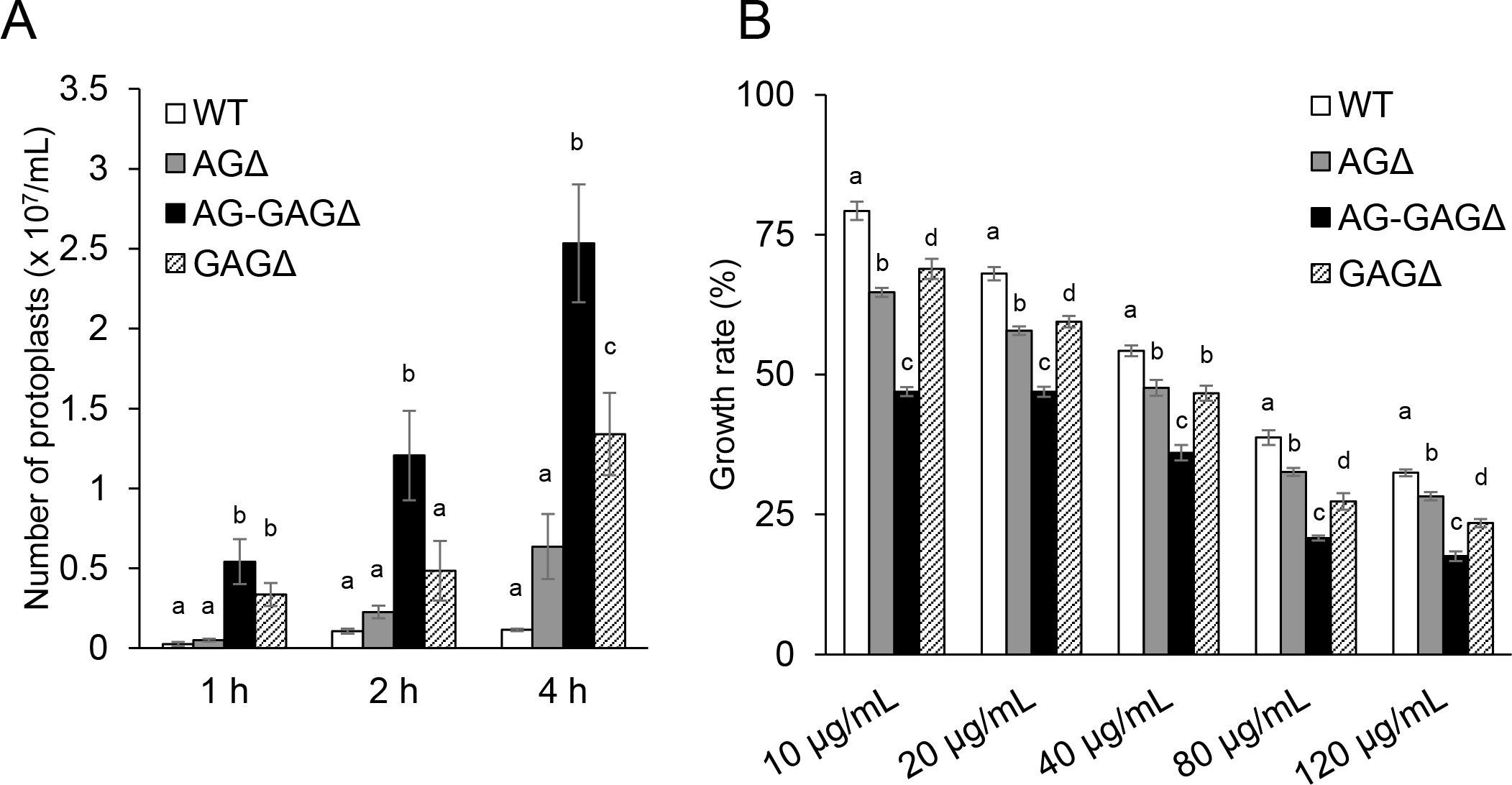
Sensitivity to Congo Red and Lysing Enzymes. (A) Mycelia cultured for 1 day were suspended in sodium phosphate buffer (10 mM, pH 6.0) containing 0.8 M NaCl and 10 mg/mL Lysing Enzymes. After 1, 2, and 4 h, protoplasts were counted under a microscope. Error bars represent the standard deviation calculated from three replicates. (B) Growth rates after 3 days on CDE medium at the indicated concentrations of Congo Red. Diameter of the colonies grown on CDE medium without Congo Red was considered as 100%. Error bars represent standard deviations calculated from three replicates. In both panels, different letters indicate significant differences within each condition by Tukey’s test (*p* < 0.05).

### Disruption of the *sphZ* and *ugeZ* genes decreases the amount of galactosaminogalactan in the cell wall

Gravelat et al. (16) quantified the GAG content as the amount of GalN after complete hydrolysis of ethanol-precipitated supernatant from *A. fumigatus* culture. To apply this approach to *A. oryzae*, we analyzed the hydrolyzed HW fractions of each strain by HPAEC. The HW fractions from both the wild-type and AGΔ strains contained 0.2–0.3 mg/g biomass GalN (Fig. 4), whereas GalN was hardly detectable in the HW fractions from the GAGΔ and AG-GAGΔ strains (Fig. 4). These results show that ugeZ and/or sphZ are essential for GAG production in *A. oryzae*.

**FIG 4.**
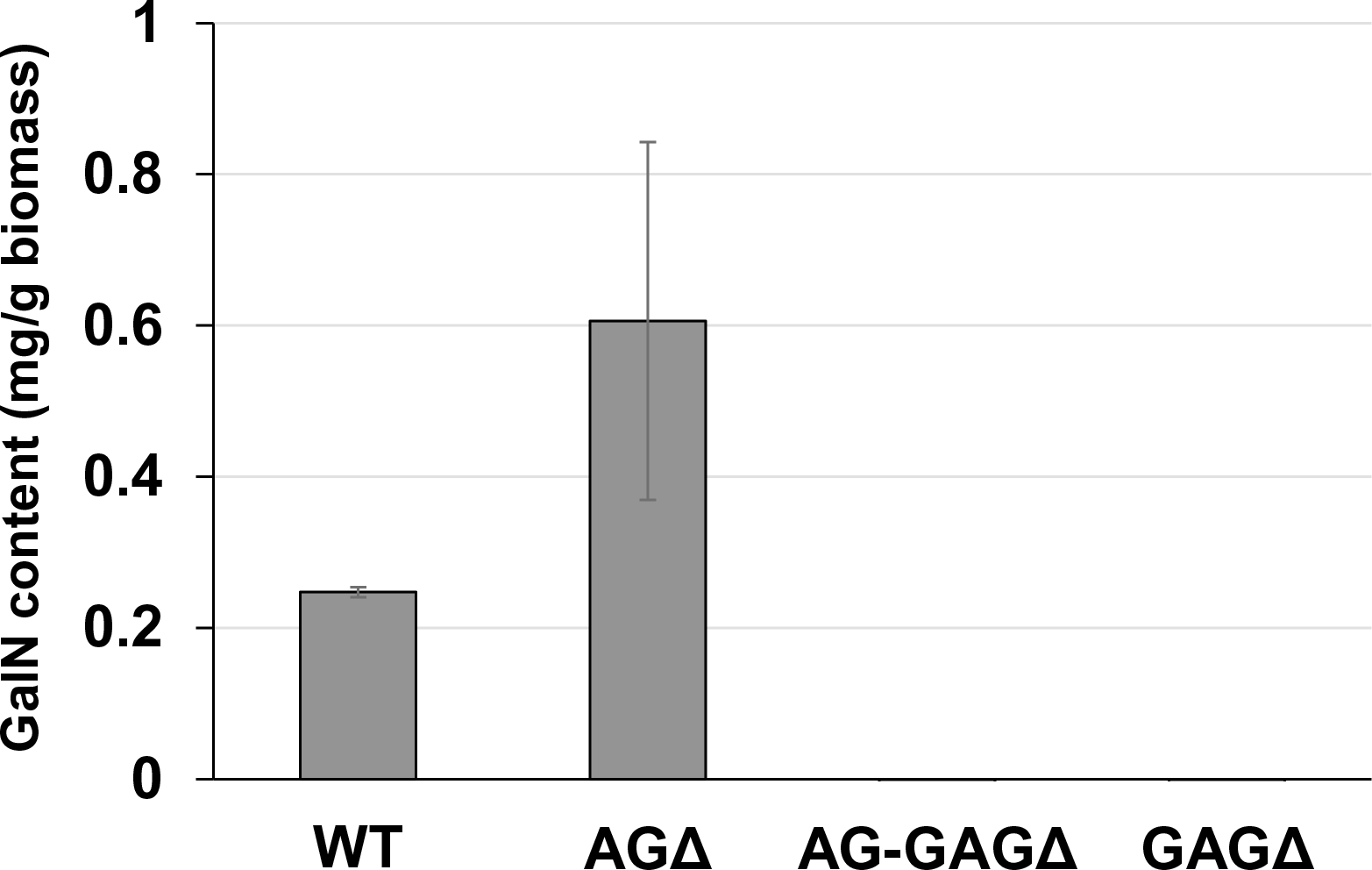
Galactosamine (GalN) content in the hot water–soluble fraction of the cell wall from the wild-type (WT), Δ*agsA*Δ*agsB*Δ*agsC* (AGΔ), Δ*agsA*Δ*agsB*Δ*agsC*Δ*sphZ*Δ*ugeZ* (AG-GAGΔ), and Δ*sphZ*Δ*ugeZ* (GAGΔ) strains. Error bars represent standard error of the mean calculated from three replicates.

### Temporally and spatially different contributions of α-1,3-glucan and GAG to hyphal aggregation in liquid culture

The complete dispersion of the AG-GAGΔ hyphae demonstrated that both α-1,3-glucan and GAG function as adhesive factors for hyphal aggregation in *A. oryzae*, and consequently the hyphae expressing both polysaccharides form pellets (Fig. 2). To analyze the temporal and spatial contribution of the two polysaccharides, wild type, AGΔ, GAGΔ, and AG-GAGΔ conidia were cultured in 48-well plates, and formation of hyphal pellets was examined (Fig. 5). The presence of α-1,3-glucan and GAG on the surfaces of conidia and germinating hyphae in liquid culture was analyzed by fluorescence microscopy with AGBD-GFP, which binds specifically to α-1,3-glucan, and lectin SBA, which binds specifically to GalNAc (Fig. 6). At the initiation of culture (0 h), conidia of all strains formed scarce aggregates (Fig. 5). Fluorescence of AGBD-GFP was observed on wild-type and GAGΔ conidia, but not on AGΔ or AG-GAGΔ conidia (Fig. 6A). SBA fluorescence was undetectable on conidia of any strains (Fig. 6A). At 3 h after inoculation, hyphae of germinated conidia of the wild-type and GAGΔ strains aggregated and formed small pellets, but aggregates of AGΔ and AG-GAGΔ germinated candida were scarce (Fig. 5). At 3 h, AGBD-GFP fluorescence was detectable on wild-type and GAGΔ germinated conidia, but none of the strains was stained with SBA (Fig. 6B). At 6 h, the wild type, AGΔ, and GAGΔ formed hyphal pellets, but aggregates of AG-GAGΔ were scarce (Fig. 5). AGBD-GFP fluorescence was observed on hyphae of the wild type and GAGΔ, and that of SBA was observed in the wild-type and AGΔ strains (Fig. 6C). At 9 h, the wild-type, GAGΔ, and AGΔ strains formed hyphal pellets (Fig. 5) similar to those formed after 24 h of culture in YPD medium. The fluorescence profiles of AGBD-GFP and SBA for each strain were similar to those observed at 6 h (Fig. 6D). The AG-GAGΔ strain hardly formed any hyphal pellets at any time point (Figs. 5 and 6). Neither conidia nor hyphae of AG-GAGΔ were stained by AGBD-GFP or SBA (Fig. 6). GFP fluorescence in the wild-type and GAGΔ strains seemed to be weaker at 0 and 3 h than at 6 and 9 h; this might have been caused by the presence of a hydrophobin layer covering the α-1,3-glucan layer (Fig. S5) (25, 26). These results indicate that hyphal aggregation caused by α-1,3-glucan was initiated just after inoculation, whereas GAG-dependent hyphal aggregation started 3—6 h after inoculation.

**FIG 5.**
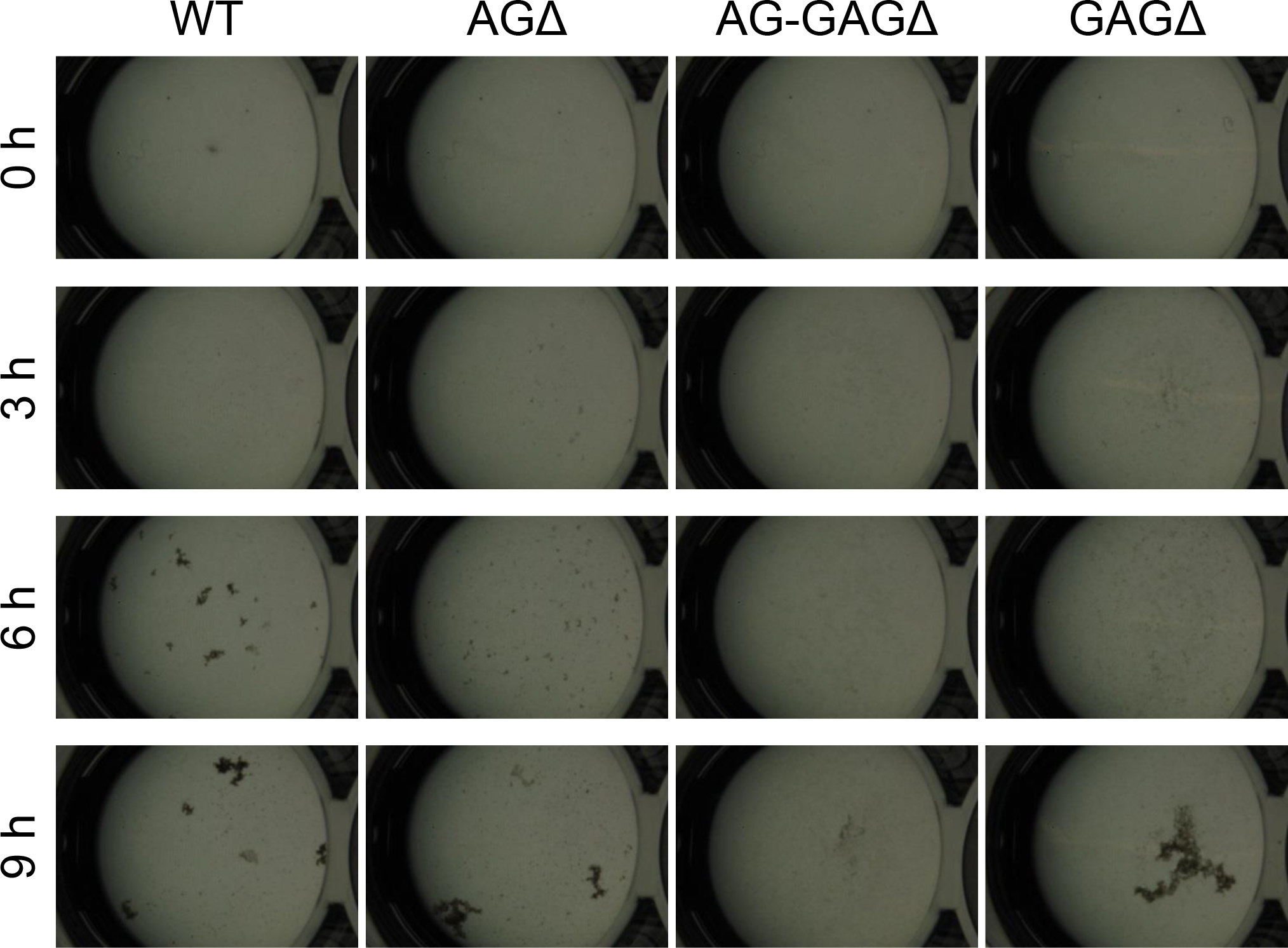
Conidial aggregation assay. Conidia (5 × 10^5^) of the wild-type (WT), Δ*agsA*Δ*agsB*Δ*agsC* (AGΔ), Δ*agsA*Δ*agsB*Δ*agsC*Δ*sphZ*Δ*ugeZ* (AG-GAGΔ), and Δ*sphZ*Δ*ugeZ* (GAGΔ) strains were inoculated into 500 µL of CDE liquid medium and incubated at 30°C with shaking (1200 rpm). Photographs were taken at the indicated time points under a stereomicroscope (magnification, ×8).

**FIG 6.**
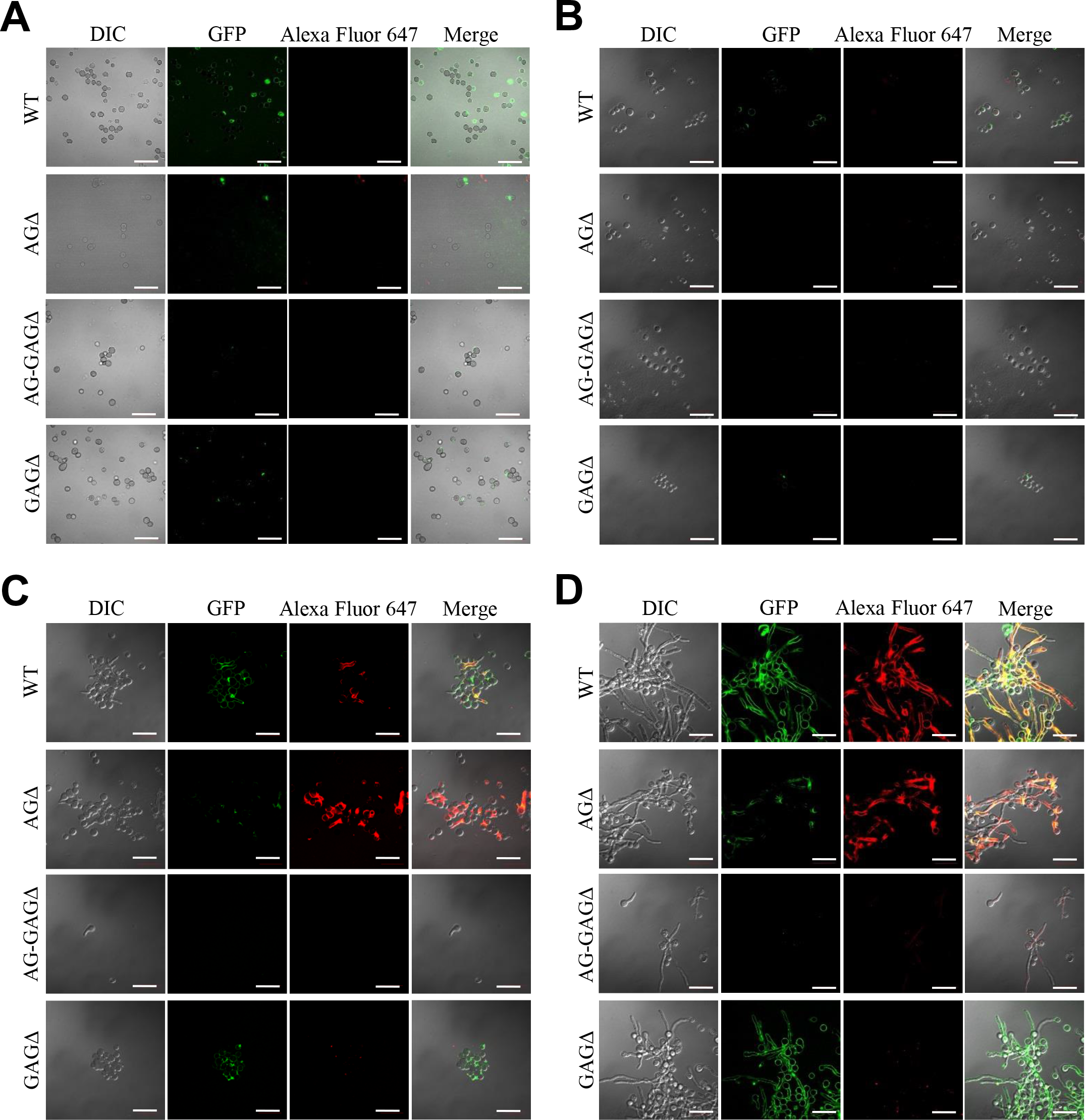
Visualization of AG and GAG in the cell wall by staining with AGBD-GFP and lectin (soybean agglutinin). Conidia (5.0 × 10^5^) of the wild-type (WT), Δ*agsA*Δ*agsB*Δ*agsC* (AGΔ), Δ*agsA*Δ*agsB*Δ*agsC*Δ*sphZ*Δ*ugeZ* (AG-GAGΔ), and Δ*sphZ*Δ*ugeZ* (GAGΔ) strains were inoculated into 500 µL of CDE liquid medium and incubated at 30°C for (A) 0, (B) 3, (C) 6, and (D) 9 h with shaking (1200 rpm). At each time point, the cells were dropped on a glass slide, fixed with 4% (w/v) paraformaldehyde, stained with AGBD-GFP and soybean agglutinin–Alexa Fluor 647 conjugate (100 µg/mL each), and observed under a confocal laser-scanning microscope (×1000). Scale bars, 20 µm.

### GAG-dependent aggregation of hyphae *in vitro* and its pH dependence

According to the previously described GAG purification method (14), we obtained the ethanol precipitates from the AGΔ strain and washed them with 150 mM sodium chloride. However, the precipitates were fully solubilized in 150 mM sodium chloride. Therefore, we developed an EtOH fractional precipitation method to isolate GAG from culture supernatants, and we obtained six fractions. The 0, 0.5, 1, 2, and 2.5 vol. fractions from the AGΔ strain contained approximately 8% of Gal and 5% of mannose, with a small amount of GalN (Fig. 7A). The 1.5 vol. fraction from the AGΔ strain contained 16% of GalN, 17% of Gal, and 4% of mannose (Fig. 7A); thus, this fraction but not the other fractions appeared to contain mainly GAG and galactomannan. The 1.5 vol. fraction from the AG-GAGΔ strain contained no GalN but contained 8% of Gal and 5% of mannose (Fig. 7A). As calculated from the composition of the 1.5 vol. fraction from the AG-GAGΔ strain, the 1.5 vol. fraction from AGΔ appeared to contain approximately 70% of GAG. To evaluate whether the aggregation of hyphae could be reproduced *in vitro*, the fractions were added to the mycelia of the AG-GAGΔ strain and mycelial aggregation was examined. Only the 1.5 vol. fraction from the AGΔ strain induced aggregation (Fig. 7B). The aggregates were stained with an SBA–Alexa Fluor 647 conjugate (Fig. 7C). The 1.5 vol. fraction from AGΔ did not form aggregates without mycelia (Fig. 7D).

**FIG 7.**
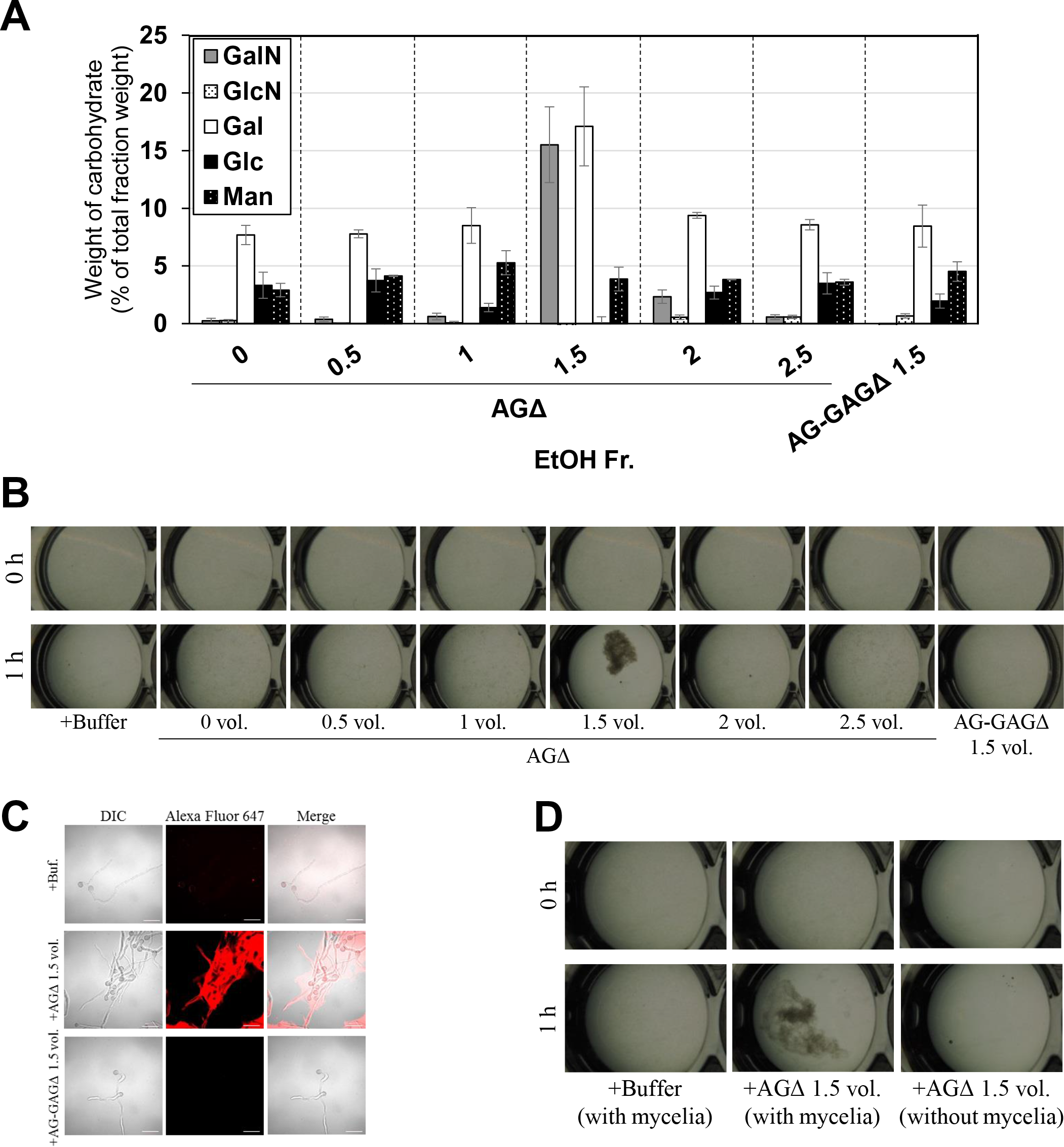
Aggregation of mycelia of the AG-GAGΔ strain induced by ethanol-precipitated GAG. (A) Composition of the fractions obtained by ethanol precipitation. (B) Mycelial suspension of the AG-GAGΔ strain (25 µL) was added into a mixture of 400 µL of water, 50 µL of 1 M sodium phosphate buffer (pH 7.0), and 25 µL of 2 mg/mL of the fractions prepared from the AGΔ or AG-GAGΔ strains, as indicated. Samples were incubated at 30°C for 1 h with shaking and examined under a stereomicroscope (magnification, ×8). (C) Mycelia incubated for 1 h in the presence of EtOH-precipitated GAG were stained with soybean agglutinin–Alexa Flour 647 conjugates and observed under a confocal laser-scanning microscope (×1000). Scale bars, 20 µm. (D) Aggregation assay with the 1.5 vol. fraction from AGΔ was performed as in (A), with or without mycelial suspension of AG-GAGΔ.

In *A. fumigatus*, GalNAc moieties in GAG are partly deacetylated and consequently positively charged (14), and we wondered whether GAG-dependent aggregation depends on pH. Addition of GAG to mycelia of the AG-GAGΔ strain led to aggregation at pH 6, 7; aggregates were scarce at pH 4, 5, and 8 (Fig. 8A). The conidia remained dispersed upon the addition of the mock fraction at any pH (Fig. 8B). These results suggest that the increased positive charge in GAG at acidic pH leads to electric repulsion among GAG chains and consequently prevents GAG-dependent mycelial aggregation. Around the neutral pH, the positive charge might be lower and consequently GAG might contribute to hyphal adhesion via non-electrostatic interactions. The reason for the absence of aggregate formation at pH 8 is unknown.

**FIG 8.**
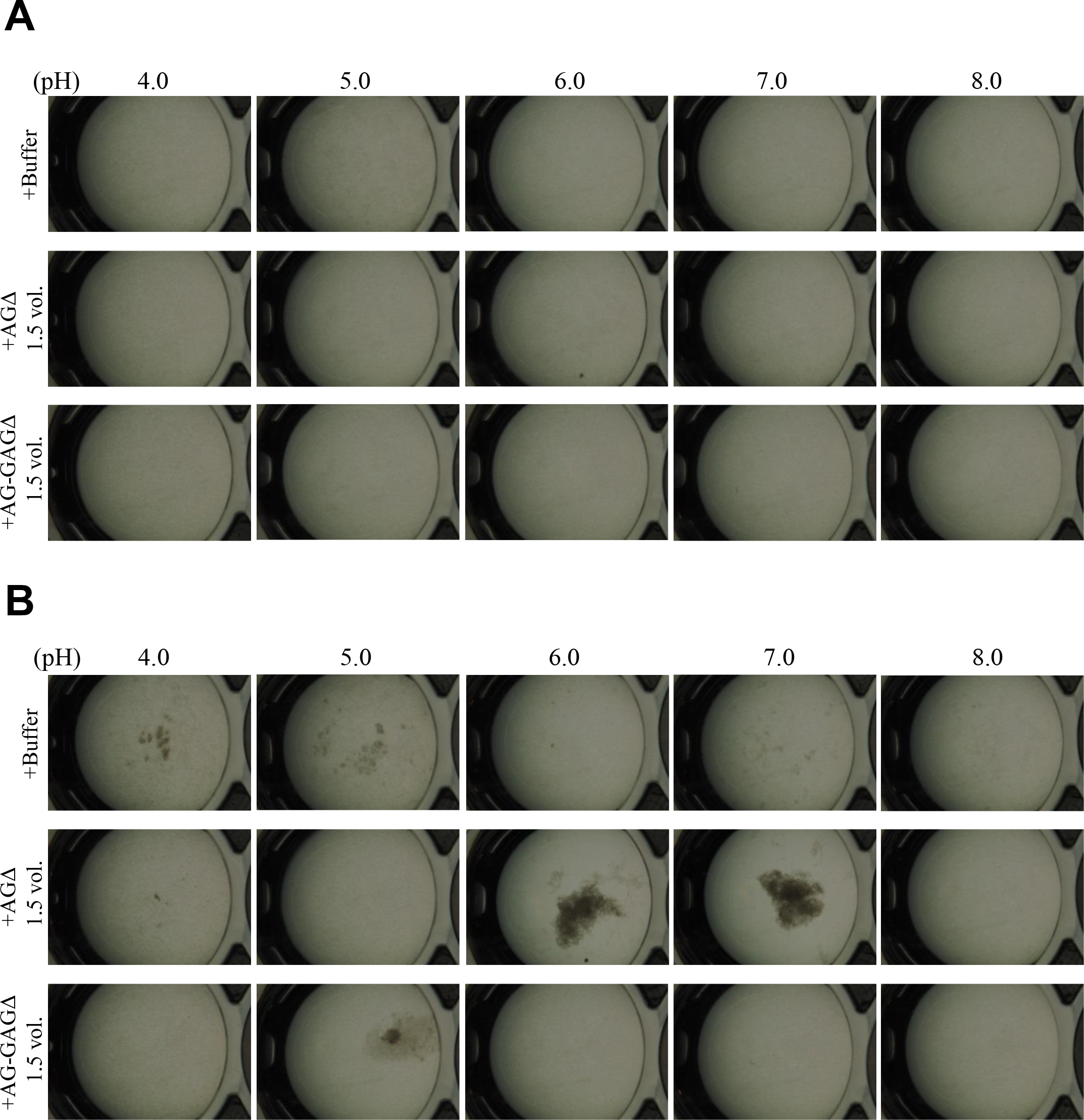
pH-dependence of GAG aggregation. Mycelial suspension of the AG-GAGΔ strain (25 µL) was added to 450 µL of buffers with different pH and 25 µL of the 1.5 vol. EtOH fraction prepared from the AGΔ or AG-GAGΔ strain as indicated. Samples were incubated at 30°C for (**A**) 0 h and (**B**) 1 h with shaking and examined under a stereomicroscope (magnification, ×8).

### GAG-dependent aggregation is caused by hydrogen bonding between polysaccharides

We hypothesized that GAG-dependent aggregation was caused by hydrogen bonding via the amino groups of GalN. To test this hypothesis, we treated GalN with or without acetic anhydrate and then evaluated the aggregation. In the presence of non-acetylated GAG, the AG-GAGΔ mycelia aggregated, similar to the results in Figure 7A, whereas GAG acetylation weakened mycelial aggregation (Fig. 9A). These results suggest that the amino groups of GalN are involved in GAG-dependent aggregation.

**FIG 9.**
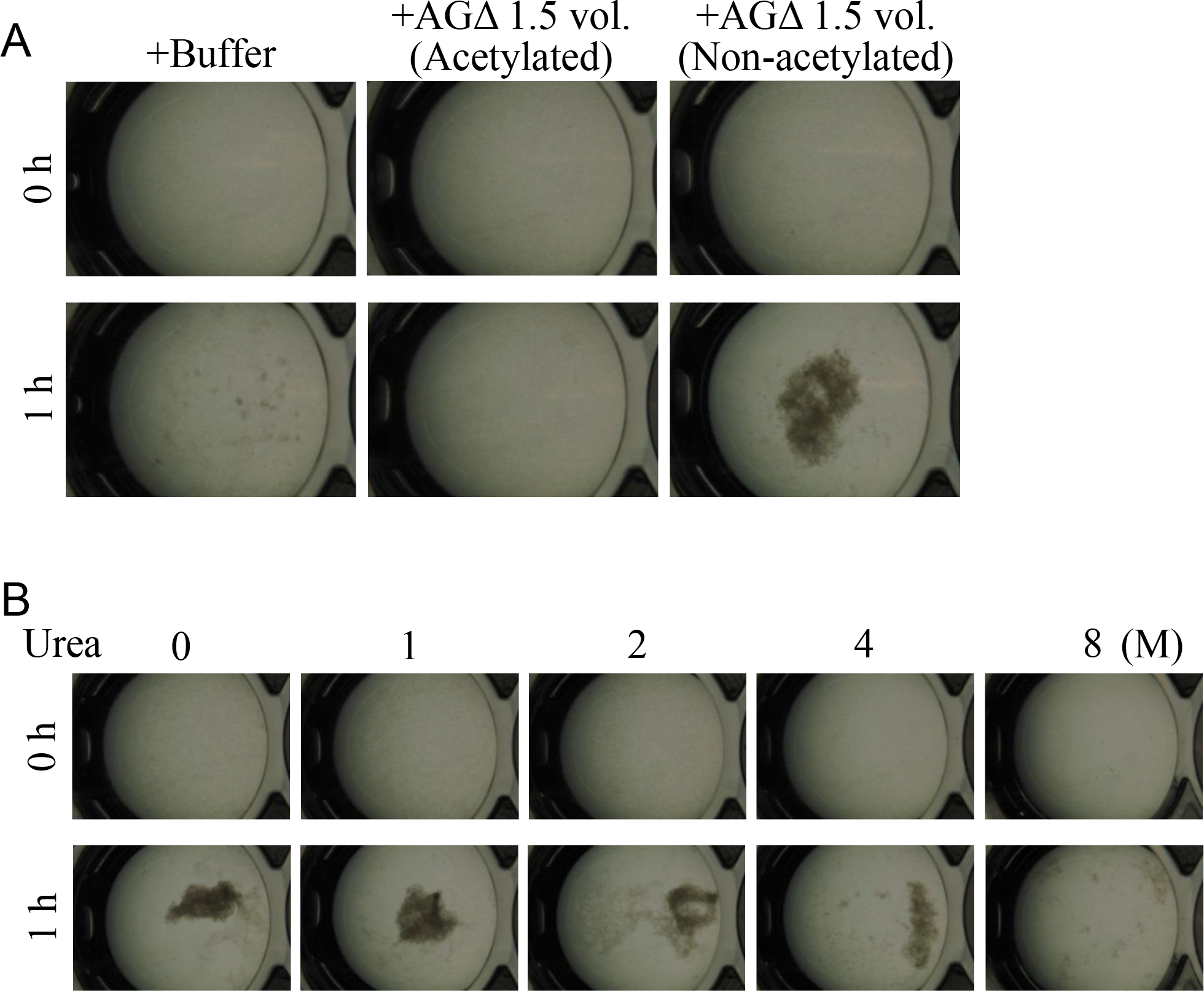
Mycelial aggregation in the presence of (A) acetylated GAG or (B) urea. (A) The amino groups of ethanol-precipitated GAG were acetylated with acetic anhydrate. Mycelial suspension of the AG-GAGΔ strain (25 µL) was added to a mixture of 450 µL of 100 mM sodium phosphate buffer (pH 7.0) and 25 µL of the 1.5 vol. EtOH fraction prepared from AGΔ (acetylated or not). (B) Mycelial suspension of the AG-GAGΔ strain (25 µL) was added to a mixture of 450 µL of 100 mM sodium phosphate buffer (pH 7.0) containing 0, 1, 2, 4, or 8 M urea, and 25 µL of the 1.5 vol. EtOH fraction prepared from the AGΔ strain. Samples were incubated at 30°C for 1 h with shaking and examined under a stereomicroscope (magnification, ×8).

To confirm that GAG-dependent aggregation relies on hydrogen bonds, we performed mycelial aggregation assay in the presence of urea, which breaks hydrogen bonds. Mycelia aggregated without urea, but aggregation was weakened by increasing urea concentrations (Fig. 9B). Taken together, these results strongly suggest that hydrogen bond formation via the amino groups of GalN is important for GAG-dependent aggregation.

### Production of a recombinant enzyme is increased in the AG-GAGΔ strain

We investigated whether hyphal dispersion would increase biomass and enzyme production in *A. oryzae*. As expected, hyphae of the AG-GAGΔ strain cultured in YPM medium were fully dispersed and those of the AGΔ strain formed smaller pellets than those of the wild-type strain (Fig. 10A). After 24 h of culture, both biomass and cutinase production were higher in the AG-GAGΔ strain (approximately 10 times) and in the AGΔ strain (4 times) than in the wild type (Fig. 10B, C). This result suggests that hyphal dispersion caused by a loss of the hyphal aggregation factors α-1,3-glucan and GAG can increase biomass and recombinant enzyme production in filamentous fungi that have α-1,3-glucan or GAG or both.

**FIG 10.**
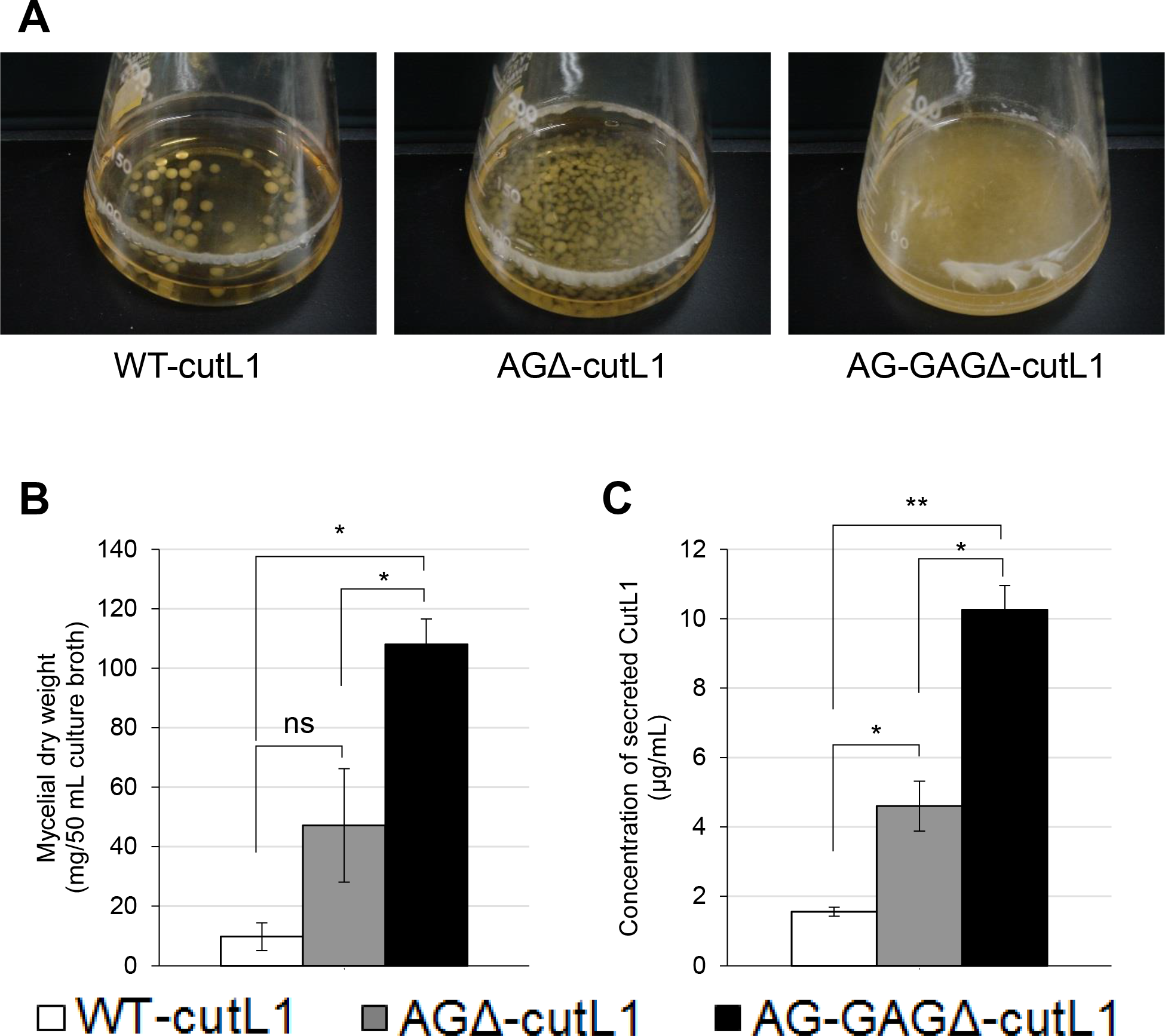
Recombinant CutL1 production by the WT-cutL1, AGΔ-cutL1, and AG-GAGΔ-cutL1 strains in liquid culture. (A) WT-cutL1, AGΔ-cutL1, and AG-GAGΔ-cutL1 strains. Conidia (final concentration, 1 × 10^4^/mL) of each strain were inoculated into YPM medium and rotated at 100 rpm at 30°C for 24 h. (B) Mycelial dry weight of each strain. Mycelia grown for 24 h were collected by filtration through Miracloth, dried at 70°C and weighed. (C) Concentration of secreted CutL1 in culture supernatants. In (B) and (C), error bars represent the standard error of the mean calculated from three replicates (\**p* < 0.05; \*\**p* < 0.01). ns, not significant.

## Discussion

Hyphae of filamentous fungi generally form large aggregated pellets in liquid culture, thus limiting the fermentative production of commercially valuable enzymes and metabolites (3, 27). Aggregation of hyphae seems to be related to their cell surface properties (7, 12, 28), but the mechanism of hyphal aggregation is not well understood. We previously demonstrated that α-1,3-glucan in the cell wall has a role in hyphal adhesion in *A. nidulans* (11, 23), and that the hyphae of α-1,3-glucan-deficient mutants of *A. oryzae* form smaller pellets than those of the wild type but are not dispersed (13). We concluded that α-1,3-glucan is an adhesive factor for *A. oryzae* hyphae, but another factor involved in hyphal adhesion remains in the AGΔ strain. Here, we focused on GAG, a component of the extracellular matrix, as a candidate adhesive factor. Lee et al. (17) revealed that GAG biosynthesis is controlled by five clustered genes (*gtb3*, *agd3*, *ega3*, *sph3*, and *uge3*) in *A*. *fumigatus*, and that similar gene clusters are conserved in various fungi, such as *A*. *niger* and *A*. *nidulans*. We found that the gene cluster is also conserved in the genome of *A*. *oryzae*.

α-1,3-Glucan contributes to hyphal and mycelial adhesion in *A. nidulans*, *A. oryzae*, and *A. fumigatus* (11–13). GAG mediates hyphal adhesion to plastic, fibronectin, and epithetical cells, and its function is related to pathogenesis in *A*. *fumigatus* (16). GAG also mediates biofilm formation in plate cultures (16). However, neither the relationship between GAG and hyphal aggregation nor the phenotype of an AG-GAG double mutant (AG-GAGΔ) has been reported. Here, we constructed AG-GAGΔ and a single mutant (GAGΔ) in *A*. *oryzae* and analyzed their growth in liquid culture. The AG-GAGΔ hyphae were completely dispersed, but the GAGΔ hyphae formed pellets larger than those of the wild-type strain (Fig. 2). These results suggest that not only α-1,3-glucan but also GAG contributes to hyphal aggregation in *A*. *oryzae*.

We investigated whether α-1,3-glucan and GAG showed temporal and spatial differences in their effects on hyphal aggregation during germination and hyphal growth. Because the germ tubes of the wild-type and GAGΔ strains aggregated at 3 h after inoculation of their conidia, whereas those of AGΔ did so at 6 h, we conclude that α-1,3-glucan is present on the surface of most hyphae just after germination and acts as an adhesive factor, whereas GAG, which is secreted and presented around the hyphal tips, contributed to hyphal aggregation at 6 h after inoculation (Figs. 5 and 6). We succeeded in *in vitro* aggregation of AG-GAGΔ hyphae by adding GAG partially purified from AGΔ strains (Fig. 7). In the presence of GAG, AG-GAGΔ mycelia aggregated at pH 6 and 7, but aggregation was reduced at acidic pH (Fig. 8B).

In the GAG biosynthetic gene cluster, *agd3* encodes *N*-acetylgalactosamine deacetylase, and GalNAc molecules in GAG chains from *A. fumigatus* are partly deacetylated (14). Disruption of *agd3* in *A. fumigatus* abolishes GAG deacetylation and results in a loss of cell wall–associated GAG (17). Positively charged amino groups in deacetylated GalNAc in GAG are thought to be required for the attachment of hyphae to negatively charged surfaces (17) and likely prevent hyphal aggregation at acidic pH because of electric repulsion; these groups would be unprotonated at pH close to neutral, in particular at the putative isoelectric point of GAG. Therefore, attachment of GAG in this pH range might be attributable to hydrogen bonding between the amino groups of GalN and the OH groups of sugar moieties in glucan of the hyphal cell wall or GAG chains pre-attached to the cell wall. Addition of GAG with amino groups acetylated by acetic anhydrate hardly induced mycelial aggregation (Fig. 9A). GAG-induced mycelial aggregation was also inhibited in the presence of 8 M urea (Fig. 9B). These observations indicate that amino group acetylation abolishes hydrogen bonding between GAG and hyphal glucans or GAG pre-attached to hyphae. Hydrogen bonds might be a major force in GAG-dependent hyphal aggregation at pH close to neutral. When hyphae aggregated by addition of GAG at neutral pH were subsequently transferred to acidic buffer (pH 4), they remained aggregated (data not shown), suggesting that, once formed, the adhesion among GAG chains is resistant to acidic conditions.

Formation of hyphal pellets limits productivity in the fermentation industry that uses filamentous fungi, including *Aspergillus* species, because the inner part of the pellet is inactive (3). In *A. niger*, titanate particles are used as a scaffold for hyphal pellets to minimize their size (27). Although physical approaches are efficient, they limit the range of culture media. The AG-GAGΔ strain produced significantly larger amounts of biomass and cutinase than did the AGΔ and wild-type strains; it did not require any scaffold particles, suggesting that controlling the hyphal aggregation factors of hyphae is an innovative approach for the fermentation industry.

We demonstrated that both α-1,3-glucan and GAG on the hyphal surface contribute to the formation of hyphal pellets and are adhesive molecules. The physicochemical properties of the two polysaccharides differ. α-1,3-Glucan is a water-insoluble major cell wall polysaccharide, whereas GAG is secreted and is a water-soluble component of the extracellular matrix. Further studies are necessary to understand the molecular mechanism underlying the interactions among α-1,3-glucan and GAG chains.

## Author Contributions

KM, AY, and KA conceived and designed the experiments. AY determined the sensitivity to LE and CR. KM and MS constructed fungal mutants. FT performed the assay of CutL1 production. KM and AS performed the Southern blot analysis. SK performed the ^13^C NMR analysis. AK and SY produced AGBD-GFP. KM and TN performed fractional precipitation of GAG. KM performed most experiments and analyzed the data.

## Acknowledgments

We are grateful to Associate Professor Toshikazu Komoda (Miyagi University) for operating the NMR spectrometer. We are also grateful to Dr. Makoto Ogata (National Institute of Technology, Fukushima College) for advising the acetylation method of polysaccharides. The manuscript was edited by ELSS, Inc. (http://www.elss.co.jp/en/).

## Supplemental Figure Legends

**Figure S1. Construction of *sphZ* and *ugeZ* gene disruption strains.** (**A**) Scheme of construction of the *sphZ* and *ugeZ* gene disruption cassette. (**B**) Strategy for replacement of the disrupted *sphZ* and *ugeZ* genes with the adenine requirement marker *adeA*. Thin arrows indicate *Pst*I digestion sites near the *sphZ* and *ugeZ* locus. (**C**) Southern blot analysis of the *sphZ* and *ugeZ* locus in the wild-type (lane 1), GAGΔ (lane 2), AGΔ (lane 3), and AG-GAGΔ (lane 4) strains using the probe indicated in (**B**).

**Figure S2. Southern blot analysis of the AG-GAGΔ-cutL1 strains.** Chromosomal DNA of the control strain (lane C) and the cutL1-overexpressing strains (lanes 1–3) was digested with *Xho*I and hybridized with the probe indicated in the upper panel.

**Figure S3. Mycelial growth of the wild-type, AGΔ, AG-GAGΔ, and GAGΔ strains on CDE agar plates.** Conidia (1 × 10^4^) of each strain were inoculated at the center of a CDE agar plate and incubated at 30°C for 4 days.

**Figure S4. Characterization of α-1,3-glucan in the cell wall of the wild-type, AGΔ, AG-GAGΔ, and GAGΔ strains.** (**A**) Composition of the AS2 and AI fractions. (**B**) Expression of the *agsB* gene. (**C**) ^13^C NMR spectra of the AS2 fractions from the wild-type and GAGΔ strains.

**Figure S5. Visualization of hydrophobin in the cell wall.** Conidia (5.0 × 10^5^) of the wild-type (WT) strain were dropped on a glass slide, fixed with 4% (w/v) paraformaldehyde, stained with fluorophore-labeled antibody against RolA, and observed under a confocal laser-scanning microscope (×1000). Scale bars, 20 µm.

